# A geometric and structural approach to the analysis and design of biological circuit dynamics: a theory tailored for synthetic biology

**DOI:** 10.1101/2020.02.18.953620

**Authors:** John P. Marken, Fangzhou Xiao, Richard M. Murray

## Abstract

Much of the progress in developing our ability to successfully design genetic circuits with predictable dynamics has followed the strategy of molding biological systems to fit into conceptual frameworks used in other disciplines, most notably the engineering sciences. Because biological systems have fundamental differences from systems in these other disciplines, this approach is challenging and the insights obtained from such analyses are often not framed in a biologically-intuitive way. Here, we present a new theoretical framework for analyzing the dynamics of genetic circuits that is tailored towards the unique properties associated with biological systems and experiments. Our framework approximates a complex circuit as a set of simpler circuits, which the system can transition between by saturating its various internal components. These approximations are connected to the intrinsic structure of the system, so this representation allows the analysis of dynamics which emerge solely from the system’s structure. Using our framework, we analyze the presence of structural bistability in a leaky autoactivation motif and the presence of structural oscillations in the Repressilator.

## 1 Introduction

Since even before the dawn of synthetic biology twenty years ago, there has been much interest in the study of designing biological systems with desired dynamical behaviors. Such a pursuit requires the development of a theoretical framework that can capture the fundamental relationship between a system’s structure and its ultimate functionality, so that desired behaviors can be engineered into the system with predictable and reliable results. There has been a rich tradition of applying conceptual frameworks from a variety of fields to the study of biological systems, ranging from physics, mathematics, engineering, computer science, and even economics, for fruitful gain. However, applying such extra-disciplinary frameworks to biology always comes with the added challenge of needing to mold the biological system under study to fit the assumptions nascent in systems from the other discipline. Because biological systems have different intrinsic properties and challenges than, say, systems in computer architecture, a theoretical framework developed to address the strengths and disadvantages of the latter will not be able to analyze a biological system optimally.

This creates an issue where, in order to best make use of conceptual frameworks from other disciplines like the engineering sciences in designing biological circuits, one must first identify the intrinsic properties of biological systems and then engineer or modify them to mitigate these properties so that they can better satisfy the requirements for these conceptual frameworks. Much of the work in the past twenty years of synthetic biology has focused on designing biological parts which minimize their biological properties (high cross-talk, probabilistic behavior, emergent behaviors) and instead exhibit the properties required for traditional engineering frameworks (modularity, orthogonality, programmed behaviors). This work has progressed both through the construction of new biological parts [1, 2] as well as the extensive characterization of existing parts [3, 4], all with the goal of making the behavior of biological systems more predictable. Although these efforts have led to great success in designing systems like genetic logic gates [5], the disconnect between the nature of biological systems and the requirements for applying engineering-based theory still persists strongly in the realm of dynamic circuit behavior.

In this work we propose an alternative approach, where we develop a new conceptual framework tailored for the specific needs of building genetic circuits out of biological parts. In addition to eliminating the need to mold biological systems to mitigate their intrinsic properties, by framing the exploration and understanding of these circuits in a more naturally biological language we may facilitate the discovery of previously-unseen insights into the design principles of biological systems. To paraphrase a lesson from one of Jeremey Gunawardena’s great perspective pieces on mathematical models, the language we use necessarily limits what we can discover and understand [6].

Our framework capitalizes on two properties of biological systems. The first is the fact that many of the biomolecular process that drive genetic circuits are enzymatic, and that therefore many of these properties are saturatable. By this we mean that when a reaction’s substrate is at a sufficiently higher concentration than its enzyme or vice versa, the reaction’s rate is well-approximated by a constant. The second property is that while the rates of biomolecular reactions often vary with environmental conditions like temperature or pH, the stochiometry of these reactions show comparatively little variation. We will use the stoichiometry of its constituent reactions to represent the structure of a biomolecular process, and therefore claim that dynamics encoded into a system’s structure rather than into the values of its rate constants will be more robust to such environmental variations.

The central insight of our framework is that a complex genetic circuit can be approximated by a set of simpler circuits, where the system transitions between these approximations by saturating or unsaturating its various components. We then find that the dynamics of the original system can be reproduced from analyzing the transitions between these approximations. Because these dynamics emerge from saturations, the resulting insights into the circuit’s dynamic behavior are phrased in the language of system saturations, which we claim is a more biologically-natural language for describing these phenomenon. Furthermore, these approximations correspond to different representations of the system’s structure, as defined above, so the dynamics that emerge from these approximations are intrinsically encoded in the system’s structure alone, rather than relying on specific values of rate constants. This ensures that such dynamics, which we term structural dynamics, will be robust to fluctuations in system parameters that are not large enough to push the system into a new saturation state.

Although our framework is superficially quite similar to the Design Space Formalism framework developed my Michael Savageau’s group [7, 8], there are important differences in both the analysis process and the intention of our work that make it a distinct framework in its own right. Furthermore, our framework serves as part of a larger theoretical foundation for models of biomolecular processes currently being developed by Xiao *et al* (in preparation) [9].

In section 2 we will begin by introducing the core concepts of the analysis framework, including the idea of partitioning a system’s dynamic behavior into structural regimes based on saturations. We will then walk through the analysis procedure for a simple case study, using the leaky positive autoactivation motif as an example of structural bistability. In section 3, we will illustrate the use of a geometric interpretation of the system’s structural dynamics to gain further insights into the leaky positive autoactivation motif. In section 4, we will apply our analysis framework to the Repressilator, to show how structural dynamics can explain the presence of oscillations. Finally, we will conclude with a discussion of the framework’s applicability to the design of genetic circuits in section 5.

Throughout this work, the more technical mathematical details will be given in appendices to facilitate a smoother reading of this manuscript. Nonetheless, a familiarity with the use of Ordinary Differential Equation (ODE) models to investigate the dynamics of genetic circuits will be helpful. Those seeking a more rigorous mathematical treatment of this framework, or those who wish to gain a fuller understanding of the theory itself, should refer to Xiao *et al* (in preparation) [9].

## 2 Preliminary Concepts

### 2.1 Introduction to saturation regimes and structural regimes

The core concept in the analysis of a system’s structural dynamics is the partitioning of a system’s state space into “saturation regimes” to approximate the complex functions driving the system’s dynamics with simpler functions, which we will term “structural regimes”. These saturation regimes are a generalization of the familiar notion of saturation in biological dynamics. As an illustration, consider the standard Hill activation function, given by

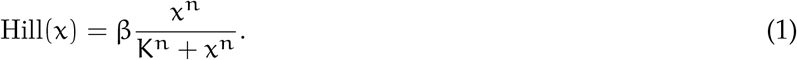

We can see that Hill(x) can be approximated by simpler functions depending on whether x is saturating the expression:

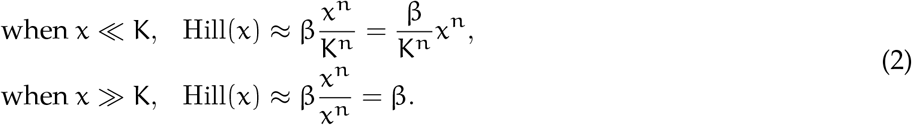

Note that each of the approximations for Hill(x) above are in monomial form, ie. they can be written as

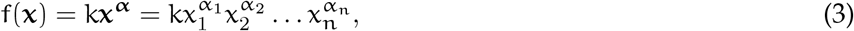

where k is some rate constant, **x** is an n-dimensional vector of system variables, and **α** is an n-dimensional vector of integers. In the case of Eq (2), when x ≪ K we have 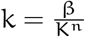 and α = n, while when x ≫ K we have k = β and α = 0.

We will say that a **saturation regime** is a window of values for the state variables **x** such that the functions of interest can be approximated by monomial forms (Eq (3)). Furthermore, we note that these monomial forms have a natural biological interpretation through the lens of mass-action kinetics. In this view, the α_*i*_ terms represent the stoichiometry of the species xi in the reaction, which corresponds to the structure of the mass-action system. Because the monomial forms in each saturation regime differ in their x_*i*_ terms, we can say that the structure of the approximation changes in different saturation regimes. Therefore we will say that a function’s specific monomial approximation in a given saturation regime is the associated **structural regime**.

Thus, in the case of the Hill activation function Hill(x), its saturation regimes are x ≪ K and x ≫ K, which correspond to the structural regimes 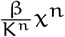 and β, respectively. This is illustrated in Fig 1a,b.

**Figure 1.**
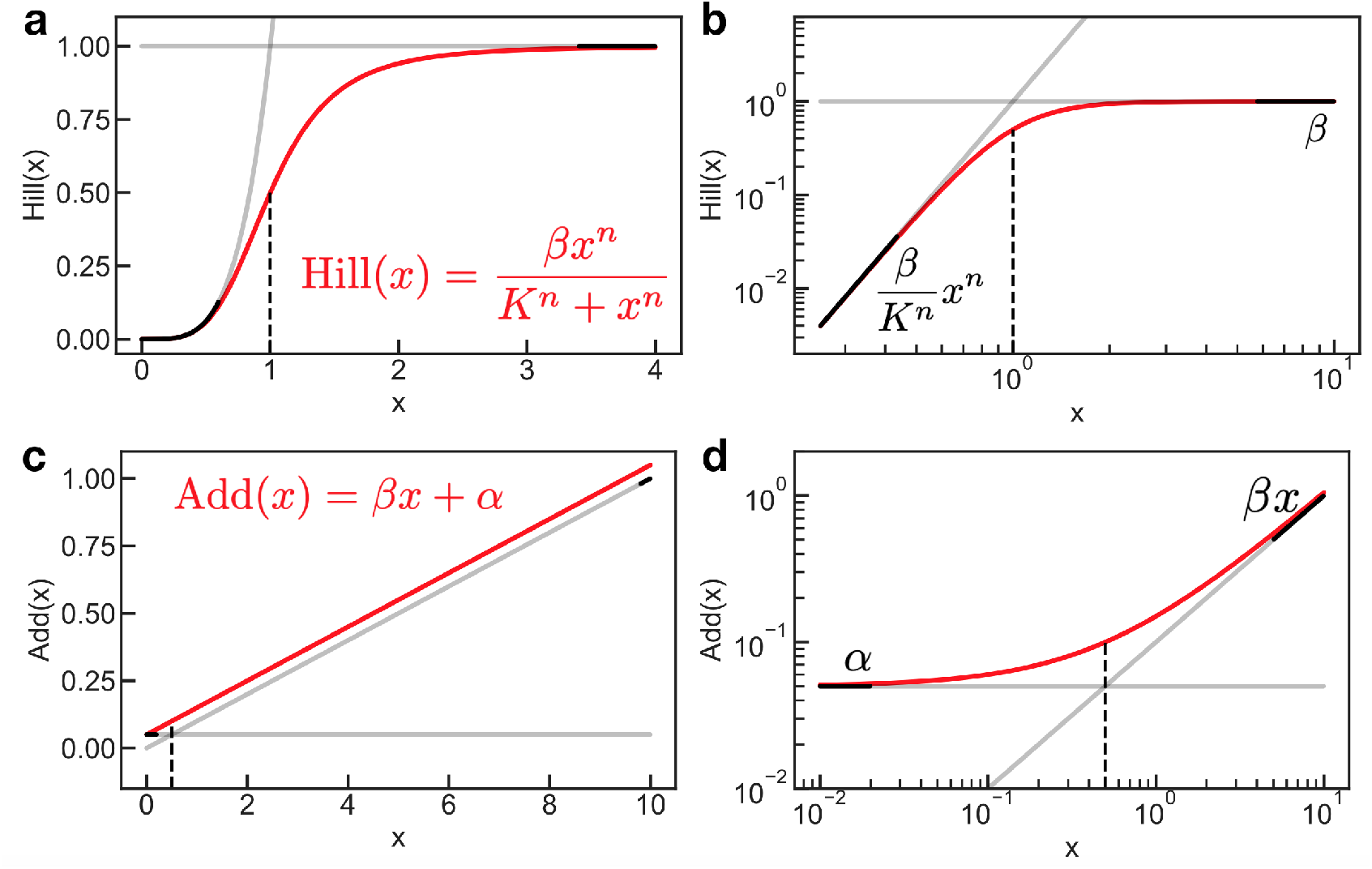
An illustration of generalized saturation and its associated regimes, applied to (**a,b**) a Hill activation function and (**c,d**) a linear addition function. The full functions are shown in red and their structural regimes are plotted in black. The structural regimes are only valid approximations of the original function within particular windows of x, which are the saturation regimes– these are schematically depicted by the boldness of the black lines. The dashed line indicates the border separating the saturation regimes– for (**a,b**) the line separates x ≪ K and x ≫ K, while for (**c,d**) the line separates x ≪ α/β and x ≫ α/β. Functions are plotted with (**a,b**) n = 4, K = 1, β = 1, and (**c,d**) β = 0.1, α = 0.05.

Furthermore, the concept of a saturation regime generalizes beyond functions which are conventionally thought of as saturatable. Consider the additive function

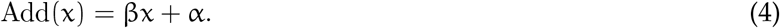

We can see that this function, too, can be partitioned into saturation regimes to approximate it with monomial structural regimes:

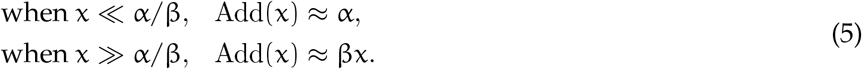

Fig 1c,d depicts the linear addition function, Eq (4), in linear space and log-log space. Note that this generalized notion of saturation for the addition function only becomes apparent in log-log space, as here the function can be seen asymptotically approaching the structural regimes, which become straight lines in the log-log plot.

Now that we’ve presented the concept of saturation regimes for individual functions, we can move on to describing how they apply to full systems. We will say that a system consists of n system variables governed by a set of n ordinary differential equations (ODEs) which can be written in the form

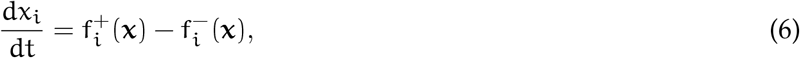

where 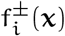 are arbitrary functions of the system variables.

We then simply approximate each individual function 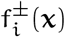 into monomial form using its saturation regimes. The system approximation will therefore take the form

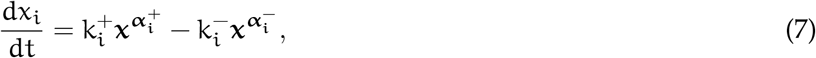

where each term above is in monomial form as described in Eq. (3).

A system that takes the form of Eq. (7) is called a **simple birth-death system**. These systems are notable because they are easy to analyze, particularly in terms of finding their fixed points. Much of the pioneering work that applied simple birth-death systems to the analysis of biochemical systems, notably the work of Michael Savageau [10], focused on this property. However, we emphasize a different property of these systems– that much of their dynamics can be determined simply from the integer exponents 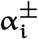, without considering the values of the rate constants 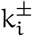. We will call these dynamics the system’s **structural dynamics**, because the mass-action view of the monomial terms in the simple birth-death system interprets the exponents 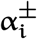 as the stoichiometry of the system’s reactions.

### 2.2 The log-derivative transformation captures structural properties of the system

In order to analyze the structural dynamics of a simple birth-death system, we want to have a procedure that will isolate the exponents **α**^±^. The log-derivative transformation h, defined for an arbitrary function f(x) as

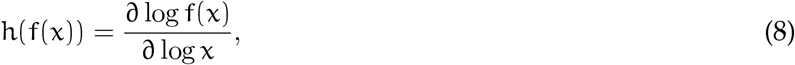

is a useful tool to do this. If f(x) is of monomial form kx^α^, for example, we get h(f(x)) = α as desired.

The generalization of the log-derivative transformation for a multidimensional system is the log-derivative matrix H, defined as

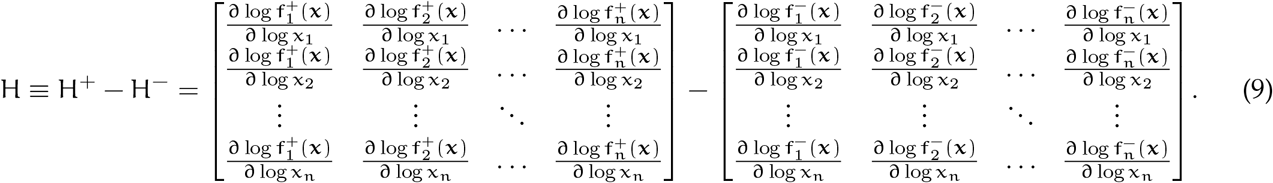

When H is applied to a simple birth-death system, the (i, j) entry of H^±^ is the integer stoichiometry for the species x_*i*_ in the birth/death process governing the dynamics of the species x_*j*_.

Furthermore, although the terms in H may look complex, in practice they are quite simple to compute– indeed, H can often be determined simply by looking at a circuit’s structure. (See Appendix B for some common log-derivative identities).

An additional reason to use the log-derivative transformation is that, at steady state, it can provide information about the system’s Jacobian. Recall that for an n-variable system governed by ODEs

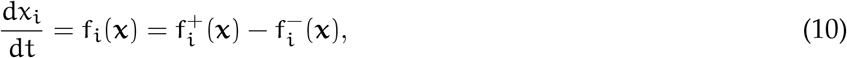

the Jacobian matrix

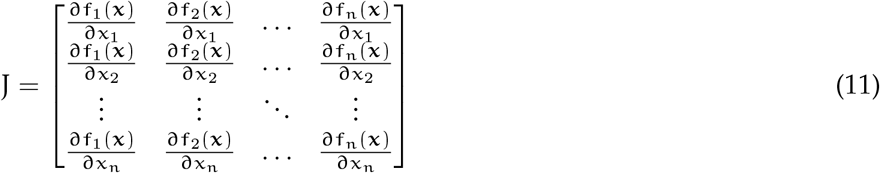

provides information about the stability of the system’s fixed points when evaluated at those points. In particular, if the Jacobian matrix is Hurwitz, i.e. its eigenvalues all have negative real parts, then the fixed point is stable.

Similarly, when the log derivative matrix H is evaluated at a fixed point, it can determine whether the point is **structurally stable**– by this we mean that the stability of the fixed point is determined solely from the system’s structure, the **α**^±^, regardless of the values of the rate constants k^±^. When certain conditions are met, we can guarantee that if H is Hurwitz then J is Hurwitz, therefore the fixed point is structurally stable. These conditions include:

1. If H and J are symmetric.
2. If H and J are triangular (ie. the circuit is a feedforward circuit).
3. If 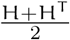 is Hurwitz.

These are not the only conditions in which H implies information about the eigenvalues of J– in general one can perform a simple test based on linear matrix inequalities to determine if H is informative (see Appendix A and [9] for details).

Note that it is possible for a fixed point to be stable but not structurally stable. In such a situation, variations in the rate parameters 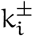 could make the fixed point unstable. Throughout this work, we will focus on situations where stability is structural, therefore it will suffice to consider H directly without computing J.

### 2.3 The analysis workflow for analyzing structural dynamics

Now that we have demonstrated the core concepts associated with the structural dynamics mindset, we will illustrate their application to the analysis of a simple circuit, the leaky positive autoactivation motif.

This circuit consits of a single gene x which is produced at some leaky rate α and degraded at some rate γ. x also activates its own production with Hill kinetics. The model for the circuit is

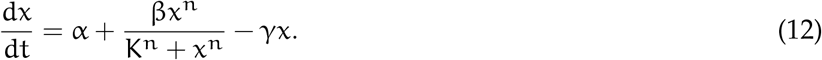

We will begin by finding the system’s saturation regimes and determining its structural regimes there– in other words, we will saturate the system’s birth and death terms until they are both monomials.

Since the death term, γx, is already a monomial, we do not need to perform any saturations there. We instead focus on the birth term, 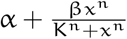. The Hill term is a good place to begin-we can see that if x ≪ K or if x ≫ K, we can approximate the system with simpler functions. In particular,

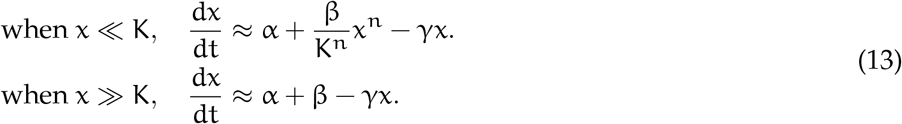

Furthermore, we can see that in the x ≪ K condition, an additional saturation exists in the additive term 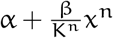.

This term is saturated when 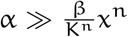, ie. when 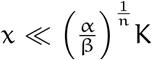. Incorporating this second saturation, we now obtain the full set of the system’s saturation regimes:

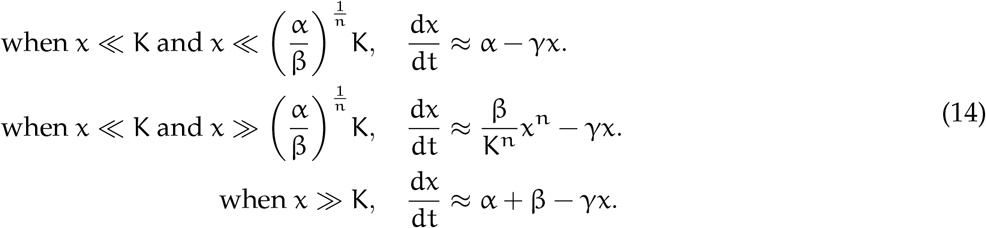

These are depicted in Fig 2.

**Figure 2.**
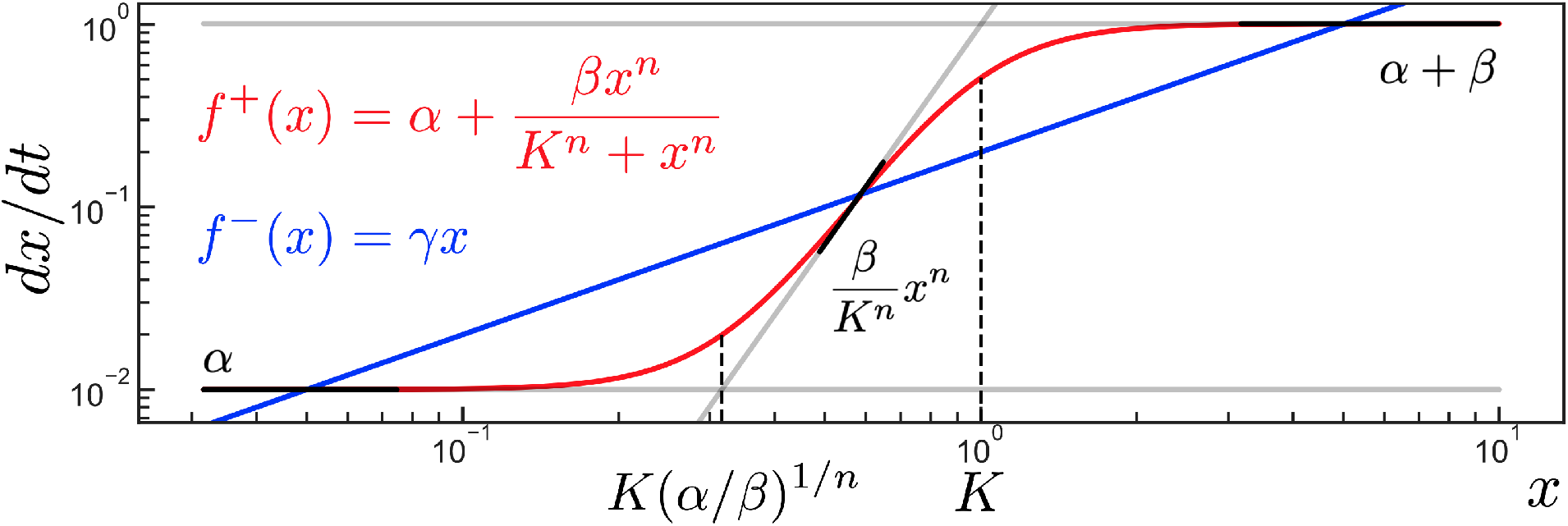
The structural regimes of the leaky positive autoregulation motif, Eq (12). The birth term and death term of the original model are plotted in red and blue, respectively. The model’s structural regimes are plotted in black, as in Fig 1.

Note that the second saturation regime in Eq (14) contradicts itself unless α ≪ β. Therefore this saturation regime can only exist when this parameter condition is satisfied, ie. when the leaky production rate is smaller than the saturated production rate from the Hill term.

The next step in our analysis is to compute the log-derivative transformation H (Eq (9)) at each structural regime to determine its structural stability. We will also set the ODEs of each structural regime to steady state in order to find the fixed points associated with each saturation regime, which we call the **saturation fixed points**.

Using the log-derivative identities given in Appendix B, we obtain

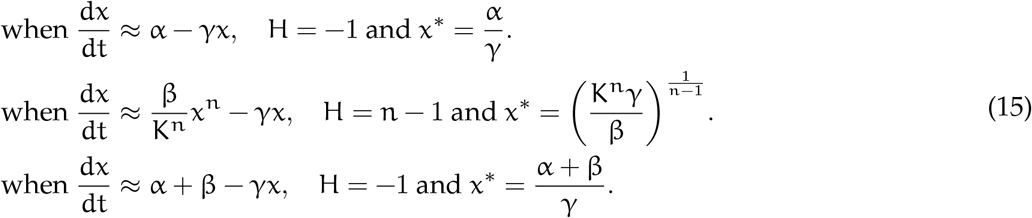

Note that if n < 1, all of the saturation fixed points in this system are structurally stable. Therefore, in order for the system to exhibit structural bistability, we require n > 1 so that one of the saturation fixed points can be unstable.

For the final step in our analysis, we determine whether the saturation fixed points actually exist within the saturation regimes they are associated with. This can be determined by applying the expressions defining the saturation fixed points to the inequalities describing the saturation regimes to get an inequality on the system’s parameters, which we call the **regime consistency conditions** because they determine whether a fixed point is consistent with its associated saturation regime.

For our system, our regime consistency conditions are

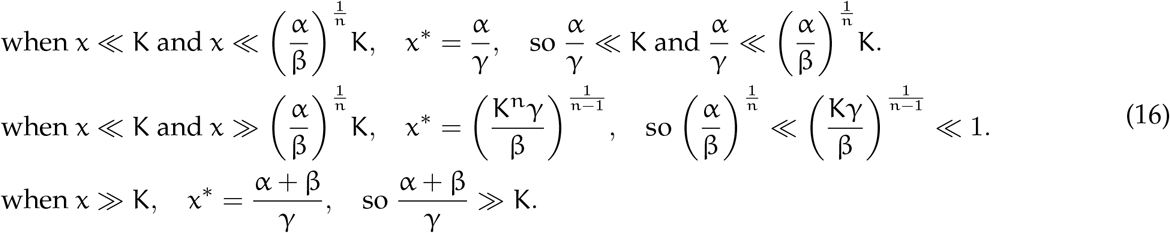

We can see that in order to satisfy each of these individual consistency conditions, the system must satisfy additional parameter constraints. For example, we see that the middle regime requires β ≫ α and β ≫ Kγ.

Now, in order to show that that the system exhibits structural bistability, we must show that none of the regime consistency conditions contradict each other, so that all three steady states can simultaneously coexist. We see that the conditions for the first two regimes above are internally consistent, as they can be written in the form

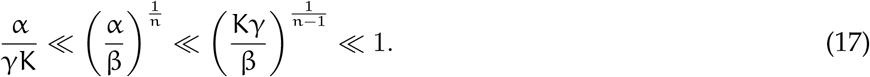

The final regime consistency condition can be written as

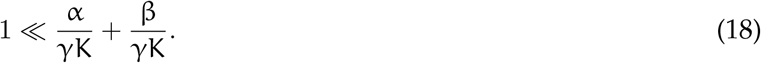

Since Eq (17) implies that β/γK ≫ 1, we can see that there is no contradiction between Eq (17) and Eq (18). Thus all three regime conditions can coexist without contradiction, so all three steady states are able to coexist as long as these parameter conditions are satisfied.

Thus we have determined that in order for the leaky positive autoactivation motif (Eq (12)) to exhibit structural bistability, we need the leaky production rate to be small compared to the activated producction rate (α ≪ β), we need cooperativity in the Hill term (n > 1), and we need strong production (β ≫ Kγ). In addition, the regime consistency conditions Eq((17) and (18)) must be satisfied. Table 1 compiles the results of our analysis.

In addition to enabling the pipeline above, the conceptual tools we have described can be used in an exploratory manner to reveal insights and intuition about a system’s dynamics. We will develop this point further in the next section.

**Table 1.** The concepts involved in the analysis of structural bistability in the leaky positive autoactivation circuit, Eq (12).

| System | $\frac{dx}{dt} = \alpha + \frac{\beta x^n}{K^n + x^n} - \gamma x$ | | |
| --- | --- | --- | --- |
| Saturation Regimes | $x \ll \left(\frac{\alpha}{\beta}\right)^{\frac{1}{n}} K \ll K$ | $\left(\frac{\alpha}{\beta}\right)^{\frac{1}{n}} K \ll x \ll K$ | $\left(\frac{\alpha}{\beta}\right)^{\frac{1}{n}} K \ll K \ll x$ |
| Structural Regimes | $\frac{dx}{dt} \approx \alpha - \gamma x$ | $\frac{dx}{dt} \approx \frac{\beta}{K^n} x^n - \gamma x$ | $\frac{dx}{dt} \approx \alpha + \beta - \gamma x$ |
| Log-Derivative Transform | $H = -1$ | $H = n - 1$ | $H = -1$ |
| Structural Stability | Stable | Stable when $n < 1$ | Stable |
| Saturation Fixed Point | $x^* = \frac{\alpha}{\gamma}$ | $x^* = \left(\frac{K^n \gamma}{\beta}\right)^{\frac{1}{n-1}}$ | $x^* = \frac{\alpha + \beta}{\gamma}$ |
| Consistency Conditions | $\frac{\alpha}{K\gamma} \ll \left(\frac{\alpha}{\beta}\right)^{\frac{1}{n}} \ll 1$ | $\left(\frac{\alpha}{\beta}\right)^{\frac{1}{n}} \ll \left(\frac{K\gamma}{\beta}\right)^{\frac{1}{n-1}} \ll 1$ | $1 \ll \frac{\alpha}{\gamma K} + \frac{\beta}{\gamma K}$ |

## 3 A geometric interpretation of structural regimes

In this section we will present a geometric interpretation of structural regimes that can give additional insights about the relationship between the system’s structure and its dynamics. The core concept of the geometric interpretation is to add variables to the system so that each possible saturation is associated with a single variable. This allows the system to be plotted in geometric space in such a way that each dimension corresponds to a unique saturation. We will continue to use the leaky positive autoactivation motif to illustrate these concepts.

### 3.1 The system expansion technique

Recall that the model for our system is

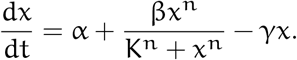

Note that this system has two possible saturations: one inside the Hill term (as in Eq (2)) and one associated with the leak term (as in Eq (5)). The goal of the geometric interpretation is to associate each saturation with a single dimension in a geometric space, so that their contribution to the system’s structural dynamics can be better visualized. We will do this by defining additional system variables until each variable is associated with one saturation. In this case, we have

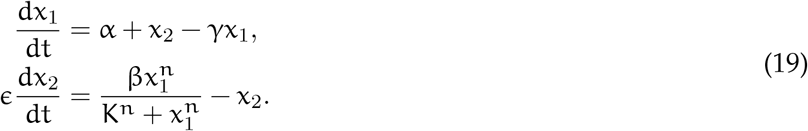

The inclusion of the ϵ ≪ 1 parameter ensures that after a very short timescale, the 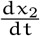 equation will equilibrate, setting 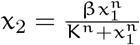. Once this occurs, then the dynamics of the expanded model (Eq (19)) will be equivalent to those of the original model (Eq (12)).

The system expansion technique can be described, in general, in the following way: suppose the original model can be written in the form

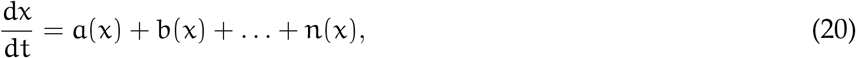

where the a(x), b(x), …, n(x) terms are arbitrary functions of the system variables. Then a valid system expansion introduces a new variable x_b_ such that

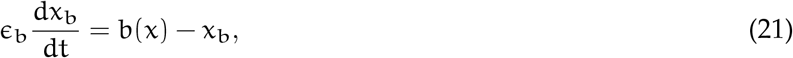

and modifies the original model to be

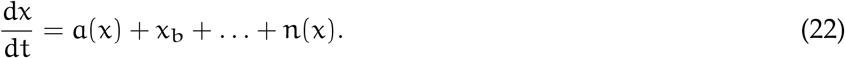

We assert ϵ_b_ ≪ 1 so that 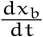 will equilibrate quickly, yielding x_b_ = b(x) so that Eq (22) will be equivalent to Eq (20) after an ϵ_b_ timescale.

For the purposes of the geometric interpretation, the functions chosen to expand into new variables should be those which are saturatable in the general sense. The procedure should then terminate once each ODE in the expanded system is associated with only one saturatable term.

### 3.2 The saturation polytope

At this point, our normal analysis procedure would have us approximate the expanded model as a simple birthdeath system using its saturation regimes. However, for the geometric interpretation, we will actually compute the log-derivative matrix H directly from Eq (19). Using the formulas given in Appendix B, we obtain

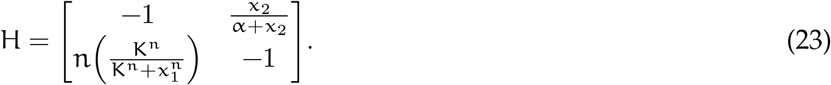

Note that each of the variable entries in H are bounded. If we write in the bounds for these entries, then we have

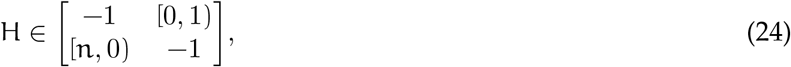

indicating that the top right entry of H ranges between 0 (when x_2_ = 0) and 1 (when x_2_ → ∞), and that the bottom left entry of H ranges between n (when x_1_ = 0) and 0 (when x_1_ → ∞).

Thus we see that H itself is bounded within a rectangle whose vertices are defined by the bounds on the variable entries of H. We will call this rectangle the **saturation polytope**, and it is visualized in Figure 3a. Because we defined our system expansion such that each variable entry in H corresponds to a unique saturation in the system, each vertex in the saturation polytope corresponds to one of the system’s saturation regimes. Furthermore, we can compute the eigenvalues of H at each vertex of the saturation polytope to determine the structural stability of the fixed point associated with each saturation regime. These are also shown in Figure 3.

**Figure 3.**
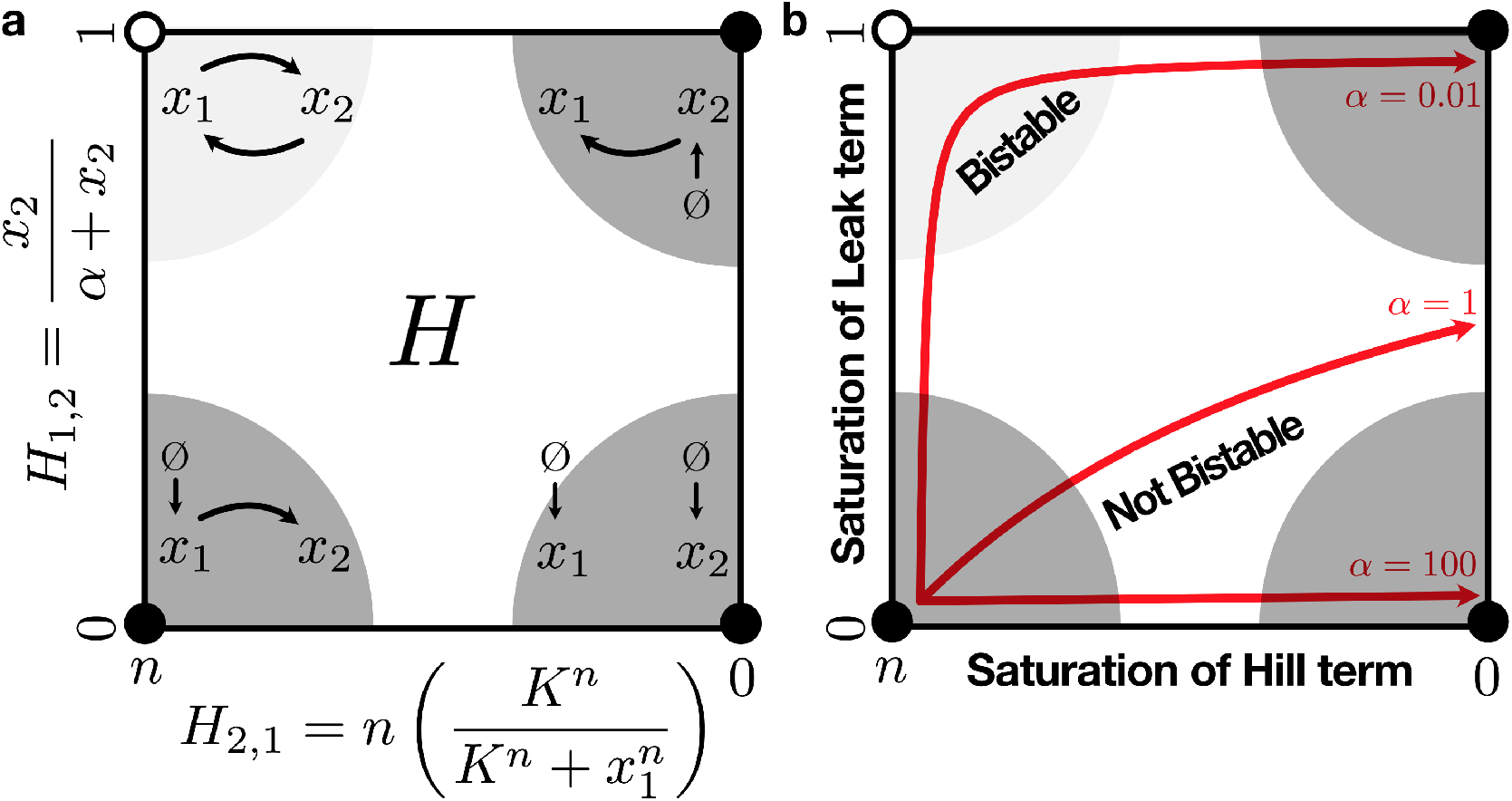
The saturation polytope for the leaky positive autoactivation motif. Filled vertices are stable and open vertices are unstable. Semicircles schematically represent the saturation regimes, ie. the regions of state space where the structural regimes are appropriate approximations of the system. (**a**) Schematic diagrams of the structural regimes. (**b**) 1-dimensional manifolds showing the trajectory of H through the polytope after dx_2_/dt has equilibrated, for different values of α. The direction of the arrows represent increasing x_1_, and all trajectories start at (n, 0) (trajectories are slightly inset from the edges of the polytope for better visualization). Manifolds are plotted with n = 4, K = 1, β = 1.

There are a number of insights that we can obtain immediately from looking at H and the saturation polytope. First, looking at the entries of H in Eq (23), we see that the only way that H can vary is via the entries that represent the system’s saturations. Because H represents the system’s structural dynamics, we conclude that *the system’s structural dynamics can only be changed by modifications in the system’s saturation state*. In particular, we see that H is completely invariant to the values of β and γ.

Second, we see that ϵ does not appear in H. By making the Hill term its own species in the expanded model (Eq (19)), we essentially introduced x_2_ as a delay term on x_1_’s autoactivation when we performed the system expansion. By setting ϵ ≪ 1, we set the timescale of the delay to be very short, but we now see that the structural dynamics are unaffected by the timescale of this delay. Therefore we see that *the structural dynamics of the system are invariant to delays in its constituent processes*. Such delays can also be implemented by introducing intermediate species into the system, and the delay invariance property will still hold (see Appendix C).

Finally, we note that after an ϵ timescale, the dx_2_/dt term in the expanded system (Eq (19)) equilibrates, yielding 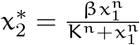. This condenses the system down to one variable, so that H can now be written as

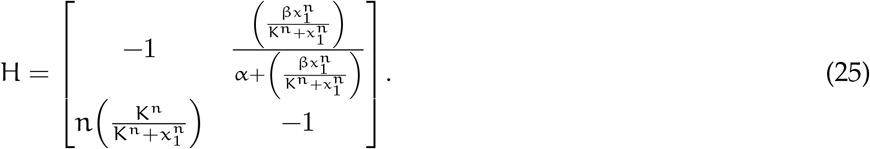

This means that as x_1_ varies, H moves in a one-dimensional trajectory through its polytope. The shape of these trajectories is determined by the parameters of the system, namely by the relationship between β and α (Fig 3b).

We can interpret these different parameterizations as the relative ease with which the system saturates its various possible saturations as x_1_ increases. In this case, the vertical axis of the polytope corresponds to the saturation of the leak term, and the horizontal axis corresponds to the saturation of the Hill term. Given that H must pass near the upper-left corner of the saturation polytope in order for the system to exhibit bistability, we thus obtain the insight that *in order for this system to exhibit structural bistability, the leak term must saturate before the Hill term*.

While conventional analyses often convey requirements for dynamical properties in the form of parameter relationships, this last point demonstrates how this geometric interpretation of structural dynamics allows insights about such requirements to emerge in a much more intuitive conceptual language. We predict that such a presentation will not only lead to a deeper understanding of the structure-function relationship in biological circuits, but will also be critical to the successful design and implementation of circuits with desired dynamical properties.

## 4 Structural oscillations in the Repressilator

Now that we have demonstrated how to analyze structural multistability in a system, we will now proceed to use the framework to analyze structural oscillations in a simple model of the well-known Repressilator circuit [11]. The key distinction between this analysis and our previous analysis of the leaky positive autoactivation motif is that here we will actively find that the regime consistency conditions are not satisfied. Instead of the system’s fixed points coexisting within their respective saturation regimes, as occurred in the case for multistability, we will instead find that each fixed point exists inside another saturation regime, such that one saturation regime will “point” to another. This pointing will create a structurally stable oscillatory cycle (Fig 4).

**Figure 4.**
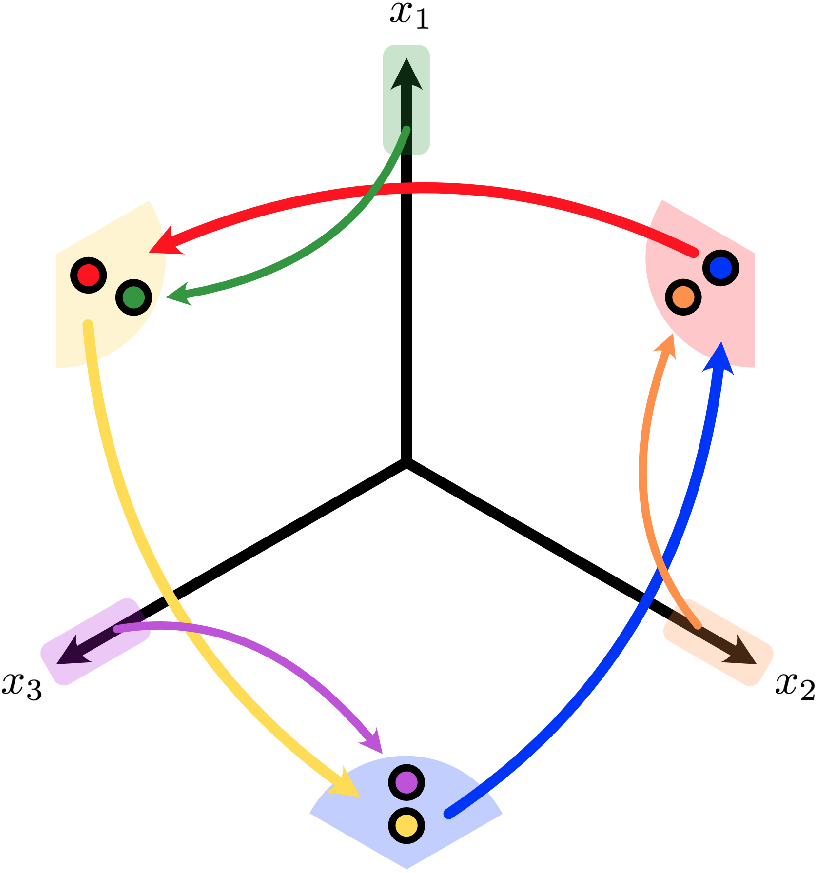
Part of the saturation polytope for the repressilator model in Eq (26), with α ≫ 1 and n ⩾ 2. When the system is in one of the saturation regimes shown above, it moves towards the stable fixed point associated with its structural regime. Because this fixed point lies in different saturation regime, once the system gets close to the point, it becomes attracted to another stable fixed point that is in a yet-again different saturation regime. This cycle forms an oscillation. The two saturation regimes corresponding to all species being low concentration and all species being high concentration are not shown here.

To begin our analysis, we will construct a simple model of the Repressilator where each component is modeled symmetrically with no leaky production, and where mRNA and protein dynamics are merged together. Then we have

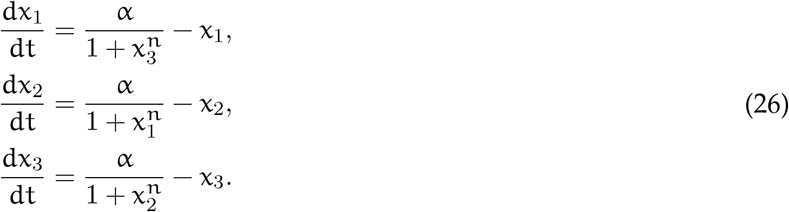

Each species x_*i*_ has two possible saturation regimes depending on the value of its repressor x_*j*_. If we specify the ordering of indices as i = 1, 2, 3 and j = 3, 1, 2, then the structural regimes for x_*i*_ (and their associated fixed points) are

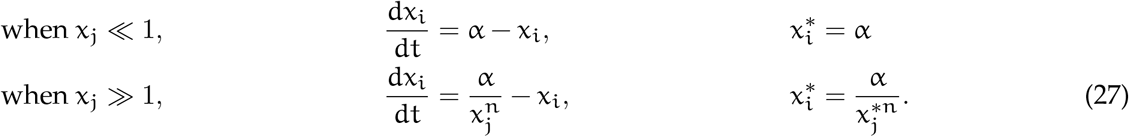

We now investigate the conditions under which this model exhibits structural oscillations. We will do this by first finding conditions for the existence of structurally stable consistent fixed points, and then negating those conditions.

Consider the fixed point in the saturation regime where x_1_, x_2_, x_3_ ≪ 1, which is 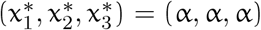. The H associated with this saturation regime is 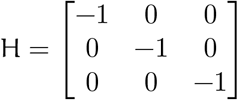, which is Hurwitz, so the fixed point is structurally stable. In order for this fixed point to exist inside the saturation regime, ie. 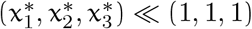, it must be the case that α ≪ 1. Therefore, in order to make this structurally stable fixed point inconsistent with its saturation regime, we must ensure α ≫ 1. Doing so ensures that when the system is in the x_1_, x_2_, x_3_ ≪ 1 saturation regime, it will tend towards a stable attractor that lies outside of the regime. Thus the system will stay in this saturation regime for only a transient period of time.

Now consider the fixed point in the saturation regime where x_1_, x_2_, x_3_ ≫ 1, which is 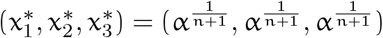. Since we have already claimed α ≫ 1, this fixed point can exist inside the saturation regime without contradictions.

However, we note that the H (same as its Jacobian) associated with this saturation regime is 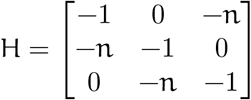 which has eigenvalues —(n + 1) and 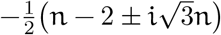. Therefore in order for this fixed point to be stable, we require n < 2. Thus if we assert n ⩾ 2, then this fixed point is no longer stable.

Hence we now have two parameter conditions that are necessary for structural oscillations in this system: α ≫ 1 and n ⩾ 2. For the six remaining regimes, we will see that no further constraints on the parameters are needed, and these regimes contain an oscillatory cycle.

For the remaining six structural regimes, it can be easily shown that H is Hurwitz and triangular, meaning that their fixed points will be structurally stable. Thus we will analyze the location of these fixed points, to see if they lie inside their associated saturation regimes or not. For convenience, in this section we will adopt the notation of (↑, ↓, ↟) to represent the saturation regime where x_1_ ≫ 1, x_2_ ≪ 1, and x_3_ ≫ 1.

We will begin with the (↑, ↓, ↑) saturation regime. The fixed point associated with this saturation regime is 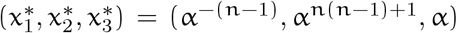, which lies in the saturation regime given by (↓, ↑, ↑). The fixed point associated with this saturation regime is 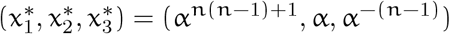, which lies in saturation regime (↑, ↑, ↓). This regime in turn has fixed point in the saturation regime we started with, (↑, ↓, ↑). This completes a cycle among the saturation regimes that proceeds indefinitely (Fig 4).

Starting at one of the three remaining saturation regimes results in convergence to this same cycle. Consider one of them, say (↑, ↓, ↓). Then the fixed point associated with this saturation regime is 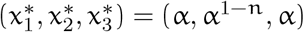. Since α ≫ 1, we see that this fixed point lies in the (↑, ↓, ↑) saturation regime, which is one of the three regimes in the cycle. The remaining two saturation regimes can be shown to converge to the cycle in the same way (Fig. 4).

We note that this conceptualization of the structural oscillations is qualitatively distinct from the conventional dynamical systems approach used in the original analysis of the Repressilator [11]. There, requirements for oscillations were determined by first determining that the system contains a single unique steady state, and then determining conditions under which that steady state becomes unstable. The oscillations are then assumed to exist in the form of a limit cycle, based on the assumption that there must be at least one stable attractor within the phase space.

In contrast, our structural mindset sees the Repressilator’s oscillations as a linked series of three stable fixed points, each of which point to each other by lying outside of their associated saturation regimes. As the system tends towards one fixed point, it will eventually enter a new saturation regime, causing the system’s structural regime to change. This invalidates the original fixed point the system was approaching, and creates a new stable fixed point (in yet another saturation regime) that the system now follows. When this loop closes, the cycle forms an oscillation. We argue that the structural mindset yields a much more intuitive conceptualization of the Repressilator’s dynamics, as it is much more closely tied to the circuit’s structure of three interlinked repressors and does not need to invoke a limit cycle to explain its oscillations.

## 5 Discussion

The case studies that we analyzed above illustrate how our theory is tailored towards the specific properties of genetic circuits as biological systems. The distinction between our theory’s conceptual framework and the more conventional dynamical systems approach to analyzing genetic circuit behavior is best highlighted in the case of the Repressilator, where instead of the conventional notion of a stable limit cycle as a singular entity we frame its oscillations as a set of stable fixed points pointing at each other. This conceptualization is much more grounded in the biological reality of the system being studied. Similarly, the insights gained from the analysis of the leaky positive autoactivation motif, by being framed in the language of saturation and structural dynamics rather than in the specific parameters of a particular model implementation, are easily generalizable to a larger class of systems beyond this specific circuit. Such generalizable insights will be essential for gaining the deep understanding of the structure-function relationship in genetic systems that is required for the predicable engineering of complex circuits.

Furthermore, our analysis approach is algorithmic, in that the same procedure is applied to analyze any circuit: First, the system is saturated until all its structural regimes are found. Then, the saturation fixed points are calculated, along with their stability. Finally, the regime consistency conditions are used to determine the location of these fixed points. Each individual step is simple to perform and understand, regardless of the complexity of the circuit being analyzed. Furthermore, although our saturation procedure is distinct from the recasting proceduredeveloped by Voit and Savageau [12]for converting generalized mass-action systems into S-System form, it is still similar in many respects, and the successful computerized automation of the recasting procedure [13, 14, 15] bodes well for the ability to eventually automate our own saturation procedure.

The algorithmic nature of our analysis contrasts with conventional dynamical systems approaches to genetic circuit analysis, which focus on phase diagram representations of the system’s dynamics– these often require numerical solution of the ODEs and a complex intuition about how changes in parameters will affect changes in objects in the phase plane. The end result is a system that does not scale well to analyzing large, complex circuits– each such circuit must essentially be analyzed from scratch with an *ad hoc* approach.

Despite the advantages of our approach, there are still a number of points which need to be developed. The main drawback is the lack of rigorous mathematical proofs and justifications for the various approximations we make, and how the dynamics of these approximations (i.e. the structural regimes) connect back to the original system. In particular, the structural regimes are good approximations for the original system when the system is far from the border of a saturation regime, but closer to this border the approximation breaks down. Currently we assume that the system dynamics transition smoothly from one structural regime to another, but this must be proven rigorously. Furthermore, the utility of the structural regimes themselves must still be expanded further– continued investigation into the connection between H and the Jacobian matrix J are ongoing.

When using this framework to aid in the design of circuits, however, this lack of rigor becomes less of an issue because the predictions of our theory can be assessed directly by experiment without relying on mathematical guarantees. Furthermore, the fact that the theory frames the dynamical properties directly through structural concepts also aids in the experimental interrogation of these systems– when designing circuits, it is often much easier to discretely add or remove structural components than it is to predictably tune arbitrary continuous parameters.

Finally, we believe that the geometric interpretation of the system dynamics enabled by our framework will be of great use in better developing an intuition of the link between system structure and function. This mindset could, for example, be incorporated into courses introducing biology students to dynamical systems analysis to better facilitate the bridge between intuitive structural ideas and their mathematical representations.

In conclusion, we hope that the advantages of this framework will not only prove useful to synthetic biologists in designing more complex circuits, but will also inspire more groups to follow the strategy of developing conceptual languages that are tailored to the needs of biological systems rather than relying on the conceptual language of other disciplines. Such an approach, when taken in complement with existing efforts to make biological systems more amenable to analysis by theoretical tools from the engineering sciences, will hopefully lead to a more holistic and fundamental understanding of the design principles governing biological systems.

## 6 Acknowledgments

J.P.M. is supported by a National Science Foundation Graduate Research Fellowship, and F.X. is partially funded by the Defense Advanced Research Projects Agency (Agreement HR0011-16-2-0049 and Agreement HR0011-17-2-0008). The content of the information does not necessarily reflect the position or the policy of the Government, and no official endorsement should be inferred.

## 7 Appendix A: How the log derivative relates to the Jacobian

Here we state and explain the relationship between log derivatives and the Jacobian to help give a concrete setting to our dicussion. For details of derivation and proof, see [cite Xiao et. al.].

Starting with a system of the form Eq (6), if we linearize the system at a fixed point **x**\* where **f**(x*) = 0, we have the following relationship between the Jacobian **J** and the log derivative matrix **H**:

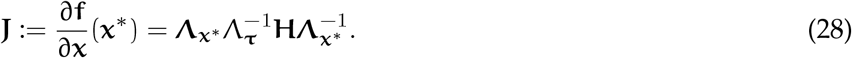

Here ***Λ***_*ν*_ denote diag(***ν***), diagonal matrix with **ν** along the diagonal. 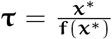 is vector of time scales, always positive. τ_*i*_ can be considered the average life time that a molecule of x_*i*_ lives before it gets degraded [16].

Note that for a simple birth-death system, ie. a system of the form Eq (7), **H** is a constant integer matrix capturing the structure, while all rate-dependence are in Λ_x*_ and Λ_τ_. So if we can show the fixed point is stable just from properties of **H**, then the fixed point is stable regardless of rates and parameters.

Inspecting the relation between Jacobian and **H** in Eq (28), immediately, we see that **J** is similar to 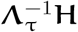, so **J**’s eigenvalues are independent of **x**\*, the position of the fixed point. In other words, if we keep **H** fixed, then stability of a fixed point depends on the time scale **τ** at that fixed point, not the location of the fixed point.

Also, directly from Eq (28), with some well-known results from linear algebra, we see **H** is Hurwitz implies **J** is Hurwitz if **H** is triangular or symmetric. For the triangular case, we know a triangular matrix’s eigenvalues are on the diagonal. Since **H** is triangular if and only if **J** is triangular, and **J**’s diagonal values are equal to **H**’s diagonal values multiplied by positive numbers from **τ** and **x**^2^, we see the sign of their diagonal values, i.e. eigenvalues, are the same. For the symmetric case, we can utilize Sylvester’s law of inertia, which states that symmetric matrices that are congruent to each other have same number of positive, zero, and negative eigenvalues. This is applicable if **H** is symmetric, as **J** is similar to 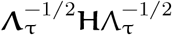, which in turn is congruent to **H**.

With some more work, utilizing the relation between Jacobian and **H** in Eq (28) allows us to state the following theorem for a test where some condition for **H** guarantees **x**\* is stable regardless of rate-dependence in **τ** and **x**\*.

### Theorem 7.1.

*The following statements are equivalent:*

1. **J** *is stable with a diagonal* **P** > 0 *certifying it, i.e*. **J**^T^**P** + **PJ** < 0.
2. **H** *is stable with a diagonal* **P** > 0 *certifying it*.
3. *There exists a positive diagonal similarity transformation of* **H**, 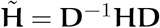 *with* **D** *diagonal with positive entries*, 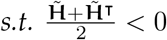.

*In particular, they imply that* **J** *is structurally stable (independent of* **x**\* *and* ***τ***).

Note that any matrix **J** is Hurwitz if and only if there exists a positive definite **P** satisfying **J**^T^**P** + **PJ** < 0. So structural stability is restricting **P** to be positive definite and diagonal [17]. The test stated in the theorem is in the form of a linear matrix inequality, which can be solved efficiently [18].

## 8 Appendix B: Some convenient log-derivative forms

For general functions f(x),

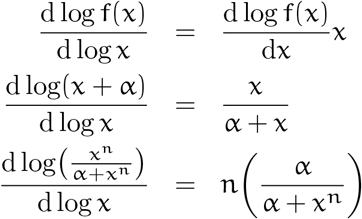

If f(x) is a monomial, ie. 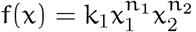, then

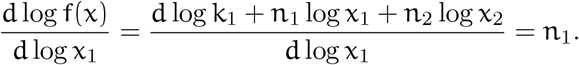

## 9 Appendix C Structural dynamics are invariant to the timescale of the system’s constituent processes

In this section we will further explore the notion of delay invariance presented in section 3.2.

Recall that we intepreted the system expansion, Eq (19), as adding a delay onto the Hill activation term in the original model. We can apply this principle further to expand the system into additional variables simply to add more delays into the system’s dynamics. Recall that our model is

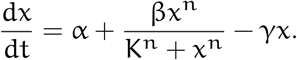

This can be expanded into

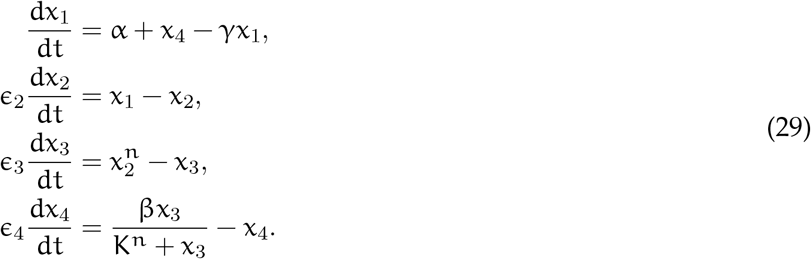

This expanded system can be interpreted as follows: the x_2_ variable now represents a distinct molecular species that is generated from x_1_ at timescale ϵ_2_– perhaps x_1_ is the mRNA and x_2_ is the protein associated with the gene X. The x_3_ variable now represents an n-mer of the x_2_ species, which takes an ϵ_3_ timescale to form. Finally, the x_4_ variable now represents the Hill activation of x_4_ activation, with a delay ϵ_4_.

If we assert that all the ϵ_*i*_ ≪ 1, then the expanded ODEs will equilibrate rapidly and the dynamics of of the expanded model will track the dynamics of the original model. However, we see that if we compute H for the expanded model, we obtain

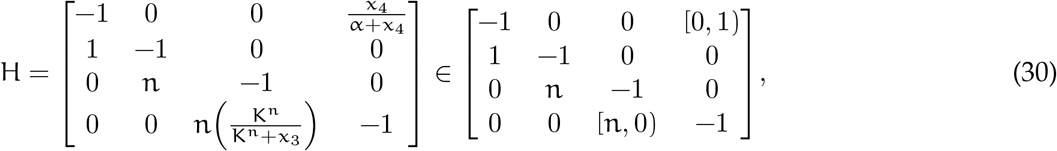

where in the right hand representation we use the same bound notation as in Eq (24).

We can see that the H matrix in Eq (30), like that in the earlier expanded H in Eq (24), can only vary via the entries which represent the system’s two possible saturations. Furthermore, if we compute the eigenvalues for H at each vertex of the saturation polytope, we find that stabilities at each fixed point are the same as in the earlier analysis. Notably, we see that once again the ϵ_*i*_ terms do not appear in H.

This means that the structural dynamics of this system are invariant not only to delays in the Hill autoactivation term, but also to translational delays and delays in multimerization. In fact, because for any function f(x) it is true that

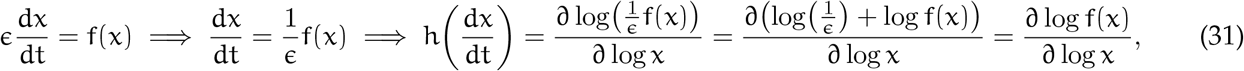

we know that the timescale terms ϵ will never appear in the log-derivative matrix H.

In fact, the implication of Eq (31) applies more generally than just to timescales– any scalar coefficient on f(x) will disappear from H in the same way. Therefore H is invariant to any scaling of the rates on its constitutent delay processes– we could have expanded the system as

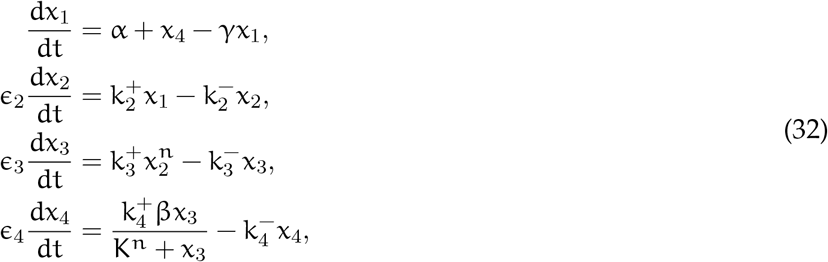

with all the parameters 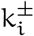 allowed to freely vary, and we would obtain the same H we obtained in Eq (30).

We can therefore conclude that *the structural dynamics of any system are invariant to the timescales of any of its constituent biomolecular processes, including those introduced by the addition of direct intermediate species*. It is important to note that this claim applies only to timescales and delays that are nascent in the original model– if we manually added new terms that altered the structure of the system, then the claim would no longer hold. The system expansion technique is able to extract these nascent delays because it does not modify the original model’s dynamics if the ϵ parameters are set to 0.

## References

[1] David L Shis, Faiza Hussain, Sarah Meinhardt, Liskin Swint-Kruse, and Matthew R Bennett. Modular, multi-input transcriptional logic gating with orthogonal laci/galr family chimeras. ACS synthetic biology, 3(9):645–651, 2014.

[2] Chen Dong, Jason Fontana, Anika Patel, James M Carothers, and Jesse G Zalatan. Synthetic crispr-cas gene activators for transcriptional reprogramming in bacteria. Nature communications, 9(1):1–11, 2018.

[3] Jacob Beal, Traci Haddock-Angelli, Markus Gershater, Kim De Mora, Meagan Lizarazo, Jim Hollenhorst, Randy Rettberg, and iGEM Interlab Study Contributors. Reproducibility of fluorescent expression from engineered biological constructs in e. coli. PloS one, 11(3):e0150182, 2016.

[4] Jacob Beal, Traci Haddock-Angelli, Geoff Baldwin, Markus Gershater, Ari Dwijayanti, Marko Storch, Kim De Mora, Meagan Lizarazo, Randy Rettberg, and with the iGEM Interlab Study Contributors. Quantification of bacterial fluorescence using independent calibrants. PloS one, 13(6):e0199432, 2018.

[5] Alec AK Nielsen, Bryan S Der, Jonghyeon Shin, Prashant Vaidyanathan, Vanya Paralanov, Elizabeth A Strychalski, David Ross, Douglas Densmore, and Christopher A Voigt. Genetic circuit design automation. Science, 352(6281):aac7341, 2016.

[6] Jeremy Gunawardena. Models in biology:’accurate descriptions of our pathetic thinking’. BMC biology, 12(1):29, 2014.

[7] Michael A Savageau, Pedro MBM Coelho, Rick A Fasani, Dean A Tolla, and Armindo Salvador. Phenotypes and tolerances in the design space of biochemical systems. Proceedings of the National Academy of Sciences, 106(16):6435–6440, 2009.

[8] Miguel A Valderrama-Gómez, Rebecca E Parales, and Michael A Savageau. Phenotype-centric modeling for elucidation of biological design principles. Journal of theoretical biology, 455:281–292, 2018.

[9] Fangzhou Xiao, Daniele Cappelletti, John Marken, John C. Doyle, and Mustafa Khammash. Structural analysis and robust design of biomolecular circuits. In Preparation.

[10] Michael A Savageau. Introduction to s-systems and the underlying power-law formalism. Mathematical and Computer Modelling, 11:546–551, 1988.

[11] Michael B Elowitz and Stanislas Leibler. A synthetic oscillatory network of transcriptional regulators. Nature, 403(6767):335–338, 2000.

[12] Michael A Savageau and Eberhard O Voit. Recasting nonlinear differential equations as s-systems: a canonical nonlinear form. Mathematical biosciences, 87(1):83–115, 1987.

[13] Rick A Fasani and Michael A Savageau. Automated construction and analysis of the design space for biochemical systems. Bioinformatics, 26(20):2601–2609, 2010.

[14] Jason G Lomnitz and Michael A Savageau. Design space toolbox v2: Automated software enabling a novel phenotype-centric modeling strategy for natural and synthetic biological systems. Frontiers in genetics, 7:118, 2016.

[15] Miguel A Valderrama-Gomez, Jason G Lomnitz, Rick A Fasani, and Michael A Savageau. Mechanistic modeling of biochemical systems without a priori parameter values using the design space toolbox v. 3.0. bioRxiv, 2020.

[16] Johan Paulsson. Models of stochastic gene expression. Physics of life reviews, 2(2):157–175, 2005.

[17] Olga Y. Kushel. Unifying matrix stability concepts with a view to applications. SIAM Review, 61:643–729, 2019.

[18] Stephen P. Boyd, Eric Feron, Venkataramanan Balakrishnan, and Laurent El Ghaoui. Linear Matrix Inequalities in System and Control Theory. SIAM, 1994.

